# Segmentation and profile-based classification of movement strategies from animal tracking data

**DOI:** 10.64898/2026.05.13.724011

**Authors:** I. Kadlec, A. Vorel, M. Sládeček, M. Kutal, M. Duľa, A. Selmovič, V. Meissner-Hylanova, N. Stier, Pešková L. Burešová, V. Barták, J. Signer

**Affiliations:** Department of Ecology, Faculty of Environmental Sciences, Czech University of Life Sciences Prague, Kamýcká129, Praha –Suchdol, 165 00, Czechia; Department of Spatial Sciences, Faculty of Environmental Sciences, Czech University of Life Sciences Prague, Kamýcká129, Praha –Suchdol, 165 00, Czechia; Department of Forest Ecology, Faculty of Forestry and Wood Technology, Mendel University in Brno, Brno, Czechia; Department of Interdisciplinary Life Sciences, Research Institute of Wildlife Ecology, University of Veterinary Medicine Vienna, Wien, Austria; Chair of Forest Zoology, Dresden University of Technology, Dresden, Germany; Wildlife Sciences, University of Goettingen, Göttingen, Germany

**Keywords:** movement ecology, movement strategy identification, Net Squared Displacement, moveprofile, profile-based classification, telemetry, trajectory segmentation, gray wolf, northern lapwing

## Abstract

1. Classifying animal movement strategies from GPS tracking data is essential for understanding space use, population dynamics and conservation planning. However, existing approaches either require strong parametric assumptions about trajectory shape, large labelled datasets (i.e. expert-annotated) for machine learning, or lack formal uncertainty quantification. These limitations create barriers for researchers working with novel species or limited sample sizes.
2. We present a profile-based classification framework consisting of three steps. First, trajectories are segmented using breakpoint detection applied to Net Squared Displacement (NSD) time series. Movement metrics are then extracted from each segment and classified by comparing them to empirically derived behavioural profiles via Z-score distances transformed to softmax probabilities. Bootstrap resampling quantifies uncertainty in the resulting classifications from both training and test data. We validated the framework through simulation experiments and applied it to GPS tracking data from two ecologically contrasting species: gray wolf (*Canis lupus*;43 individuals) and northern lapwing (*Vanellus vanellus*;15 individuals).
3. Simulations showed that 5–10 training segments per movement strategy suffice for reliable classification, with overall accuracy of 91.1%across residential, floating and dispersal strategies. Segment duration of 30–60 days was required for confident discrimination of residential and floating behaviour. For wolves, the framework clearly distinguished residency, floating or dispersal (91.2%of segments classified with >50%probability). For lapwings, migration was identified with high confidence, while residential–floating discrimination reflected genuine ecological ambiguity confirmed by domain experts, with bootstrap confidence intervals transparently flagging uncertain cases.
4. The profile-based framework provides an accessible, interpretable alternative to parametric NSD fitting and machine learning approach, requiring modest training data while delivering probabilistic classifications with honest uncertainty estimates. An R package (moveprofile) implementing the complete workflow is freely available. The framework is applicable to any tracked species where distinct movement strategies can be identified by experts’ knowledge.

## 1 Introduction

Animal movement is one of the most fundamental processes shaping ecological systems, with far-reaching consequences for individual fitness, population dynamics, and ecosystem functioning (Jeltsch et al., 2013;Nathan et al., 2008). How animals move through their environment reflects underlying movement strategies shaped by the interplay between internal state, navigation capacity, and external factors, including resource distribution, predation risk, and reproductive opportunities (Abrahms et al., 2017;Nathan et al., 2008). Among vertebrates, these strategies span a remarkable spectrum, from localised residency within stable home ranges to long-distance migration between seasonal habitats, exploratory nomadism in search of ephemeral resources, and natal dispersal that maintains genetic connectivity among populations (Bunnefeld et al., 2011;Teitelbaum & Mueller, 2019). These distinct movement strategies are not merely descriptive categories but represent induced responses to ecological pressures that directly influence survival, reproduction, and population persistence (Merkle et al., 2016;Mueller & Fagan, 2008). Understanding and identifying movement strategies of individual animals has become increasingly important for conservation planning (Sawyer et al., 2019), predicting disease transmission dynamics (Altizer et al., 2011), managing human-wildlife conflict (Linnell et al., 2020), and forecasting species responses to environmental change (Tucker et al., 2018). Yet, identifying these strategies from tracking data remains methodologically challenging.

Despite the ecological importance of movement strategies, quantifying them from tracking data remains methodologically challenging, as existing approaches each carry distinct assumptions and limitations (Bunnefeld et al., 2011;Edelhoff et al., 2016;Spitz et al., 2017). A commonly used approach fits different parametric models to observed Net Squared Displacement (NSD) of trajectories, each representing a distinct movement strategy(Bunnefeld et al., 2011;Singh et al., 2012). While NSD is clearly mechanistically interpretable, this framework requires strong assumptions about trajectory functional forms and provides limited guidance when observed patterns deviate from expected shapes (Börger & Fryxell, 2012). Alternatively, clustering approaches offer data-driven alternatives by grouping similar movement metrics without parametric assumptions, yet they struggle with arbitrary cluster number selection and provide hard classifications without probabilistic assessments of uncertainty (Edelhoff et al., 2016). Machine learning methods, including random forests and neural networks, have demonstrated high classification accuracy when trained on large, balanced datasets (Wijeyakulasuriya et al., 2020), yet their black-box nature complicates ecological interpretation, and their data requirements often exceed what is available for rare species or behaviours (Valletta et al., 2017). Critically, most existing approaches lack formal frameworks for quantifying classification uncertainty arising from limited training data, leaving practitioners without quality indicators to assess when classifications are reliable versus when additional data collection or expert validation is warranted.

Here, we present a profile-based classification framework that classifies both movement strategies and the uncertainty inherent in their identification. The method constructs reference behavioural profiles from labelled training segments (sections of trajectories with known movement strategy), capturing characteristic patterns in NSD across time. Classification proceeds by comparing new segments to these profiles, yielding not only strategy assignments but also confidence metrics derived through bootstrap resampling. This quantifies uncertainty arising from limited training data and reflects it in the reported classification probabilities (Davison & Hinkley, 1997;Efron & Tibshirani, 1994).

The approach requires only a few training segments per movement strategy from different individuals, far fewer than for most machine learning methods. This allows the method to be used for rare species or behaviours where extensive data collection is impractical or impossible. Researchers can control which movement metrics are used based on species ecology, maintaining methodological consistency while accommodating biological diversity. Importantly, classifications remain interpretable: one can identify which specific movement characteristics drove each decision, facilitating biological insight rather than black-box predictions.

Our primary objectives are threefold. First, we develop and validate the profile-based classification methodology through simulation experiments with known ground truth before applying it to empirical data, evaluating its performance across two species, a mammal (gray wolf, *Canis lupus*) and a bird (northern lapwing, *Vanellus vanellus*) with fundamentally different movement characteristics. These species exhibit contrasting movement strategies and ecological contexts. Wolves alternate between territorial residency of established packs, floating behaviour characterised by exploratory movements without permanent settlement, and short but critically needed natal dispersal. Lapwings frequently alternate between residency during incubation and chick rearing, floating, and long-distance migration between breeding and wintering grounds. Second, we implement bootstrap uncertainty quantification to distinguish high-confidence classifications from ambiguous cases warranting expert review or additional data collection. Third, we provide practical guidance on implementation of decisions including training data requirements, movement metric selection, and temporal resolution of analysis.

## 2 Materials and Methods

We divide the classification into several sequential stages (Figure 1). We start by aggregating GPS positions into time windows of several hours reflecting the actual biological question, then summarize data within time-window by calculating their centroids and calculate a time series of NSD. Next, we search for breakpoints to partition trajectories into segments, from which we extract movement metrics. After which expert-labelled training segments establish reference profiles for each behavioural class;we then classify unknown segments using Z-score distances transformed to probabilities with softmax and we quantify uncertainty by bootstrapping over segments.

**Figure 1.**
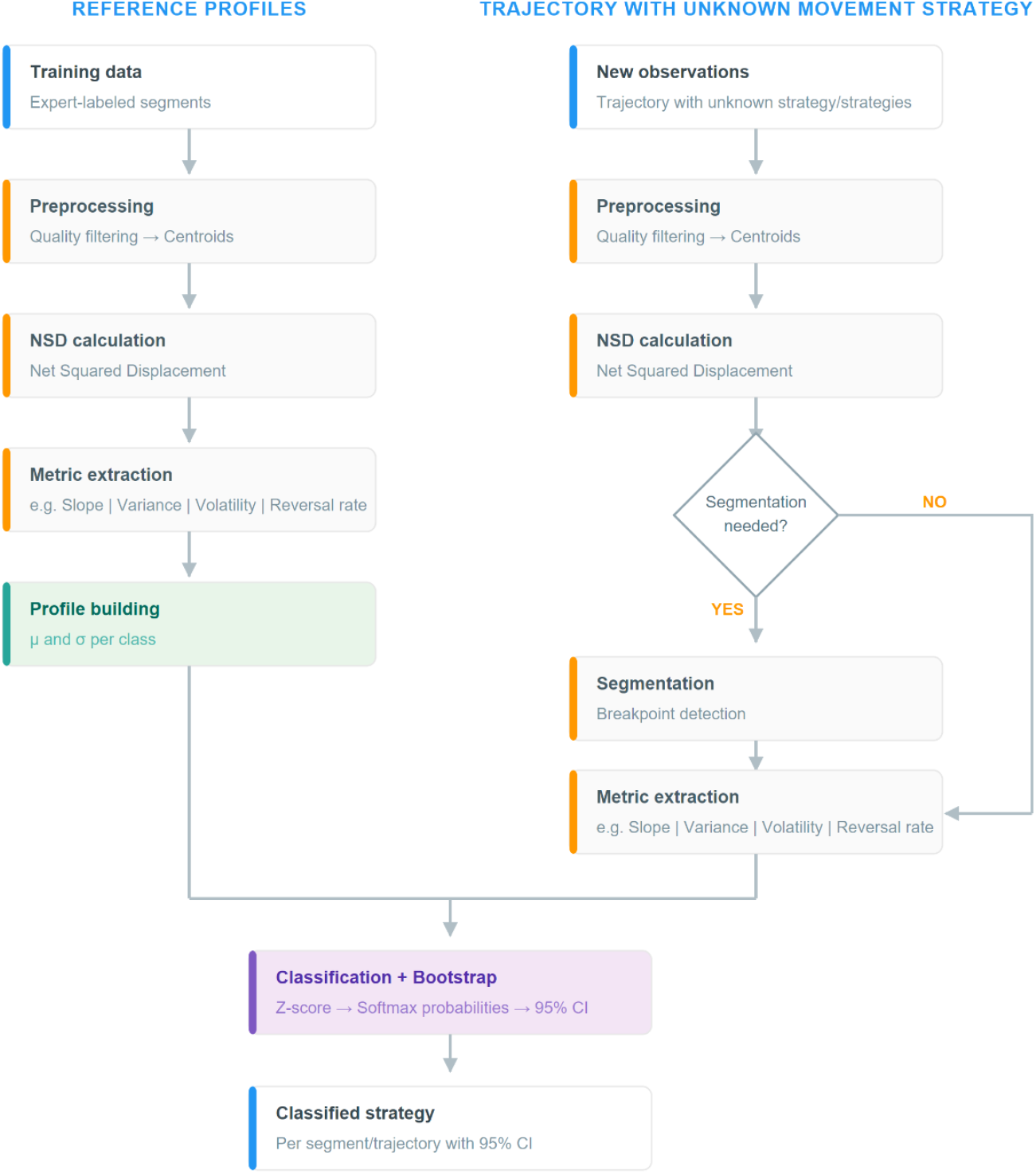
Workflow of the profile-based movement classification framework. Training data (left path) and test data (right path) undergo preprocessing, NSD calculation, and metric extraction. Training segments build reference profiles (mean and standard deviation per behavioural class), which are used to classify test segments via Z-score distances converted to softmax probabilities. Bootstrap resampling quantifies classification uncertainty, yielding probability estimates with 95%confidence intervals.

### 2.1 Data preprocessing and NSD calculation

We designed the method to work with GPS telemetry data from a tracked animal. We apply quality filters to retain only high-precision locations (e.g., dilution of precision DOP <6). To reduce noise from fine-scale movements, we aggregate raw positions into temporal centroids. For a given time interval Δt, we calculate the mean position of all GPS fixes within that interval:

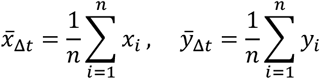

where *n* is the number of observations within the interval. This approach supports multiple temporal resolutions (e.g., 3h, 6h, 12h, 24h, …), allowing users to chose which aggregation scale use to select for desired movement strategy detection. Shorter temporal intervals capture fine-scale behavioural transitions, whereas longer intervals suppress daily variation and highlight long-term displacement patterns. The optimal temporal resolution depends on studied species biology and on intended the research question. We recommend using daily-based centroids, but in some cases, this may be too rough (see case studies).

From the centroid time series, we calculate Net Squared Displacement (NSD), a widely used metric in movement ecology that quantifies the squared Euclidean distance between an animal’s current position and its starting location (Bunnefeld et al., 2011). For a position at time *t*, we define NSD as:

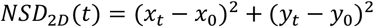

where (*x*_0_, *y*_0_) represents the initial position and (*x*_*t*_, *y*_*t*_) is the position at time *t*. NSD provides a cumulative measure of spatial displacement over time, making it sensitive to changes in movement behaviour. Unlike instantaneous metrics such as step length or turning angle, NSD integrates movement history, allowing for the detection of long-term behavioural patterns.

For species that exhibit significant vertical displacement (e.g., altitudinal migrants, marine divers), NSD can be extended to three dimensions where *z* represents altitude or depth:

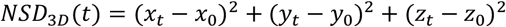

### 2.2 Trajectory segmentation

Movement strategies are not static;animals transition between different movement strategies in response to ecological cues. To identify these transitions objectively, we apply a grid-search breakpoint detection algorithm to the NSD time series. For this purpose, we evaluate all possible breakpoint combinations within user-defined temporal ranges. For each candidate breakpoint combination, we fit and assess a piecewise linear model using Akaike Information Criterion (AIC), which balances model complexity against goodness-of-fit. We select the breakpoint configuration with the lowest AIC as optimal segmentation. This approach partitions the trajectory into discrete segments, each representing a distinct movement strategy.

Within each segment j (a contiguous section of the NSD time series between two consecutive breakpoints), we model NSD as:

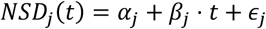

where α_j_ is the intercept, β_j_ is the regression slope, representing the rate of change in squared displacement per unit time and ε_j_ represents residual error. The slope has a direct ecological meaning: values near zero indicate residency, while positive or negative values suggest migration or exploration, respectively.

We use three criteria that control segmentation: 1) minimum gap between breakpoints (default threshold: 5 days), 2) minimum segment size (default: 5 observations), and 3) search step size (default: 3 days). Users should adjust these values based on species ecology and GPS sampling frequency. Note that uncertainty arising from segmentation parameter selection is not propagated through bootstrap resampling and represents an additional source of uncertainty in the final classifications.

### 2.3 Training data and profile construction

Classification requires a training dataset of expert-labelled segments from tracked individuals. A researcher should perform this classification with expert knowledge of the species’ecology through visual inspection of trajectories. The number and definition of movement strategies depend on the study species;for example, wolves may exhibit residential (territorial), floating (nomadic), and dispersal behaviours, while migratory birds show residency, some type of nomadism/floating and migration.

For each training segment, we calculate the movement metrics described in Section 2.4, yielding a training matrix **X** ∈ℝ^(n×p), where n is the total number of training segments across all behavioural classes and p is the number of metrics. Each row of **[X, y]** represents one segment, and each column **X** represents a metric value:

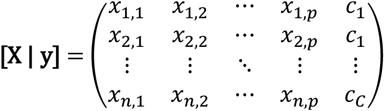

where the last column represents the label vector **y**, with c i ∈{1, …, C} denoting the behavioural class of segment i. From this matrix, class-specific mean vectors **μ**_c and standard deviation vectors **σ**_c are computed separately for each behavioural class c:

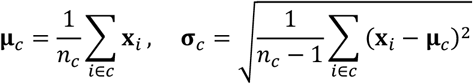

where *n*_*c*_ is the number of training segments belonging to class c. These vectors define the characteristic “profile”of each behavioural class in metric space and serve as reference profiles for classifying unknown segments.

Training data should ideally span multiple individuals and, where possible, different temporal periods to capture natural variability within each behavioural class and avoid overfitting to individual-specific movement patterns.

### 2.4 Movement metrics

From each segment, movement metrics quantify behavioural signatures. The framework is flexible regarding metric selection;users should choose metrics distinguishing between relevant movement strategies based on the ecology of the studied species.

We evaluated six candidate metrics (slope, residual standard error, rolling variance, volatility, reversal rate, and autocorrelation) and recommend the following four core metrics that collectively provided the best discriminatory performance in simulation experiments:

**Slope (β) of NSD** quantifies the rate of spatial displacement, derived from linear regression of NSD on time:

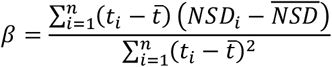

Steeply positive slopes indicate rapid, directional movement away from the origin, characteristic of dispersal or outward migration, while negative slopes may reflect a return phase or inward migration, and slopes near zero suggest residency.

**Rolling variance** captures local variability in NSD over a sliding window of width w:

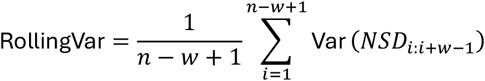

High rolling variance indicates unstable or exploratory movement patterns, while low values suggest consistent behaviour within the segment.

**Volatility** measures short-term movement consistency as the mean absolute change between consecutive NSD values:

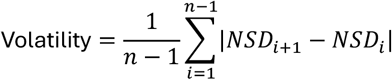

High volatility characterizes erratic, fluctuating displacement typical of floating behaviour, while low volatility indicates smooth, gradual changes in displacement characteristic of animals remaining within a confined area. Unlike rolling variance, which reflects variability across a local time window, volatility captures movement roughness at the level of individual consecutive steps, providing complementary information at a finer temporal scale.

**Reversal rate** quantifies oscillatory behaviour as the proportion of directional changes in the NSD series:

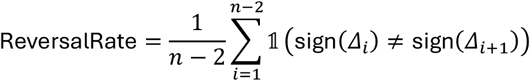

where Δ_i_ = NSD_i_+_1_ −NSD_i_. High reversal rates indicate back-and-forth movements without net displacement, while low rates characterize unidirectional movement.

These four metrics collectively capture directionality (slope), local stability (rolling variance), short-term consistency (volatility), and oscillatory patterns (reversal rate), providing effective discrimination across residency, migration, and exploratory strategies in our case studies.

Beyond the four core metrics, the framework includes additional optional metrics that may improve classification in specific ecological contexts. **Residual standard error** of the slope regression quantifies the deviation of observed NSD values from the fitted linear trend;low values indicate that movement closely follows a linear trajectory, while high values reflect irregular or mixed movement patterns. **Autocorrelation** quantifies directional persistence as the lag-1 autocorrelation of the NSD time series, measuring the correlation between NSD(t) and NSD(t−1);high values indicate persistent directional movement characteristic of dispersal, while low values reflect erratic or oscillatory displacement typical of floating or residential behaviour. Researchers may also implement custom metrics tailored to their study system, provided they show clear separation between the behavioural classes of interest and carry interpretable ecological meaning. For example, altitudinal migrants might benefit from elevation-based metrics (elevation range, vertical volatility), while marine species could incorporate depth profiles.

### 2.5 Classification and uncertainty quantification

To classify an unknown segment, we use a distance-based approach in which standardised Z-scores quantify how many standard deviations the unknown segment deviates from each reference profile. For an unknown segment with metric vector **x**_unknown_, we calculate standardized distances (Z-scores) to each behavioural class *c*:

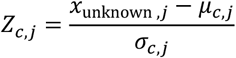

where *j* indexes movement metrics for c indexes behavioural classes. The overall distance to class *c* is the mean absolute Z-score:

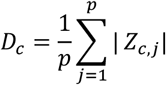

This metric quantifies how many standard deviations the test segment deviates from the typical profile of each behavioural class. Smaller distances indicate greater similarity to the class.

To convert distances into interpretable class membership probabilities, we apply the softmax transformation:

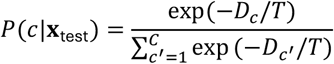

where *C* is the number of classes and *T* is a temperature parameter (set to *T* = 1 by default). The softmax function converts raw distances into class membership probabilities, assigning higher probabilities to closer profile matches while ensuring all probabilities sum to unity. The temperature parameter T controls classification sharpness: at low values, the highest-probability class dominates (approaching a hard assignment), while higher values produce more uniform distributions. At T = 1 (default), probabilities directly reflect relative distances without additional sharpening;a segment equidistant between two strategies receives similar probabilities for both, transparently flagging ambiguous cases.

To quantify classification uncertainty, we implement bootstrap resampling applied after segmentation, to the identified segments. Bootstrap resampling therefore does not propagate uncertainty arising from breakpoint detection or segmentation parameter selection, which represent additional sources of uncertainty acknowledged in the Limitations section. Profile parameters are estimated from B_profile resamples (default: 100) of the training data, each sample of the same size as the original training set and drawn with replacement, independently within each behavioural class. For each resample, class parameters (μ_c, σ_c) are recalculated and classification probabilities recomputed, capturing uncertainty from finite training sample sizes. Double bootstrap (nested resampling of both profile building and unknown segment data) additionally resamples observations within the unknown segment B_unknown times (default: 100), yielding B_unknown ×B_profile = 10,000 iterations that propagate variability from both training and unknown segment data. Resampling of training data is performed within each behavioural class independently of individual identity;multiple segments from the same individual are treated as separate observations. We report results as mean probability with 95%confidence intervals, for example: Residential: 78.2%[62.1%, 91.3%]. We interpret classification reliability based on the relationship between confidence intervals across strategies. Non-overlapping confidence intervals between classes indicate unambiguous classification. Overlapping intervals warrant additional scrutiny;however, if the confidence interval of the assigned class does not contain the mean probability of any alternative strategy, the classification remains reliable. If the confidence interval of the assigned class contains the mean probability of an alternative strategy, the classification should be treated with caution and verified by expert review.

### 2.6 Simulation design

To evaluate framework performance under controlled conditions with known ground truth, we simulated trajectories representing three movement strategies with characteristic NSD dynamics (Figure 2). We constructed residential trajectories with low baseline NSD values and multi-frequency oscillations representing daily and weekly activity patterns, with occasional short-duration excursions that return to a central location. We generated floating trajectories using a random walk process with variable step sizes, frequent directional reversals, and occasional large displacements, but without sustained directional movement. We built dispersal trajectories around a strong positive linear trend with superimposed stopovers creating temporary plateaus and autocorrelated noise reflecting movement persistence. Each trajectory type incorporated observation noise calibrated to produce realistic metric distributions comparable to empirical tracking data. Simulation code and parameters are available at GitHub repository (https://github.com/kadleci/moveprofile) as a part of package *moveprofile*.

**Figure 2.**
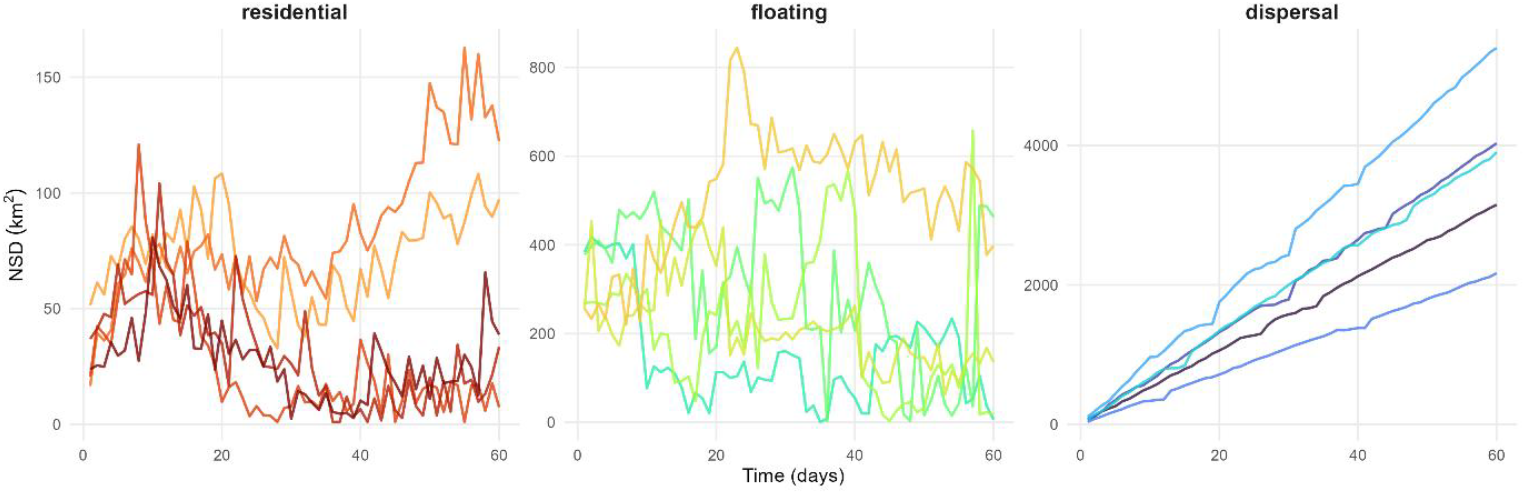
Simulated Net Squared Displacement (NSD) trajectories for three movement strategies. Residential trajectories (a) show low-amplitude oscillations with occasional excursions returning to a central location. Floating trajectories (b) display erratic, high-variance displacement without directional persistence. Dispersal trajectories (c) exhibit strong positive trends with occasional stopovers. Five replicates shown per strategy;note different y-axis scales.

### 2.7 Empirical data

We applied the classification framework to GPS tracking data from two species (gray wolf and northern lapwing) with contrasting movement ecologies.

#### Gray wolves (*Canis lupus*)

GPS telemetry data were obtained from wolves monitored across Central Europe (Czech Republic, Germany, Austria, and Poland) between 2016 and 2025. The dataset comprised 43 individuals: 13 used for training with expert-assigned behavioural classifications (18 segments total), and 30 for classification. Training and test datasets contained no overlapping individuals. Collars recorded locations at variable intervals (from 1 to 12 hours);positions were aggregated into daily (24-hour) centroids. Wolves exhibit three movement strategies: territorial residents maintaining stable home ranges, floating individuals engaging in nomadic exploration without settlement, and dispersers undertaking directed long-distance movements.

#### Northern lapwings (*Vanellus vanellus*)

GPS tracking data were collected from 15 lapwings breeding in the Czech Republic and migrating to wintering grounds in Spain and France between 2024 and 2026. Four individuals (47 segments) with expert-assigned classifications served as training data;11 individuals were classified. Positions were aggregated into 12-hour centroids to capture faster behavioural transitions. Three movement strategies were distinguished: residential phases during incubation and chick rearing, floating phases representing exploratory movement and stopovers, and migration phases involving rapid directed displacement.

## 3 Results

We evaluated the profile-based classification framework through simulation validation followed by empirical application to two species. First, we assessed classification performance using simulated trajectories with known ground truth to establish baseline accuracy and examine the effects of training sample size, segment duration, and metric selection. We then applied the framework to GPS tracking data from gray wolves (*Canis lupus*) and northern lapwings (*Vanellus vanellus*), representing contrasting movement ecologies at fundamentally different spatial and temporal scales.

### 3.1 Simulation Validation

Classification accuracy depended strongly on training sample size (Figure 3, panel A). With only two training segments per behavioral class, mean probability assigned to the true class averaged approximately 50–65%depending on the strategy, with residential performing worst (median ~48%) and dispersal best (median ~68%). This reflects the challenge of estimating profile parameters from minimal data, where sampling variance in the training set propagates directly to classification uncertainty. Performance improved substantially with additional training data: at five segments per class, dispersal and floating classifications approached 90%probability, while residential reached approximately 75%. By ten segments per class, all three strategies achieved median probabilities exceeding 85%, with dispersal and floating approaching ceiling performance. Beyond ten segments, marginal improvements were minimal, suggesting that modest training datasets of 5–10 segments per class suffice for reliable classification when behavioral categories are well-defined.

**Figure 3.**
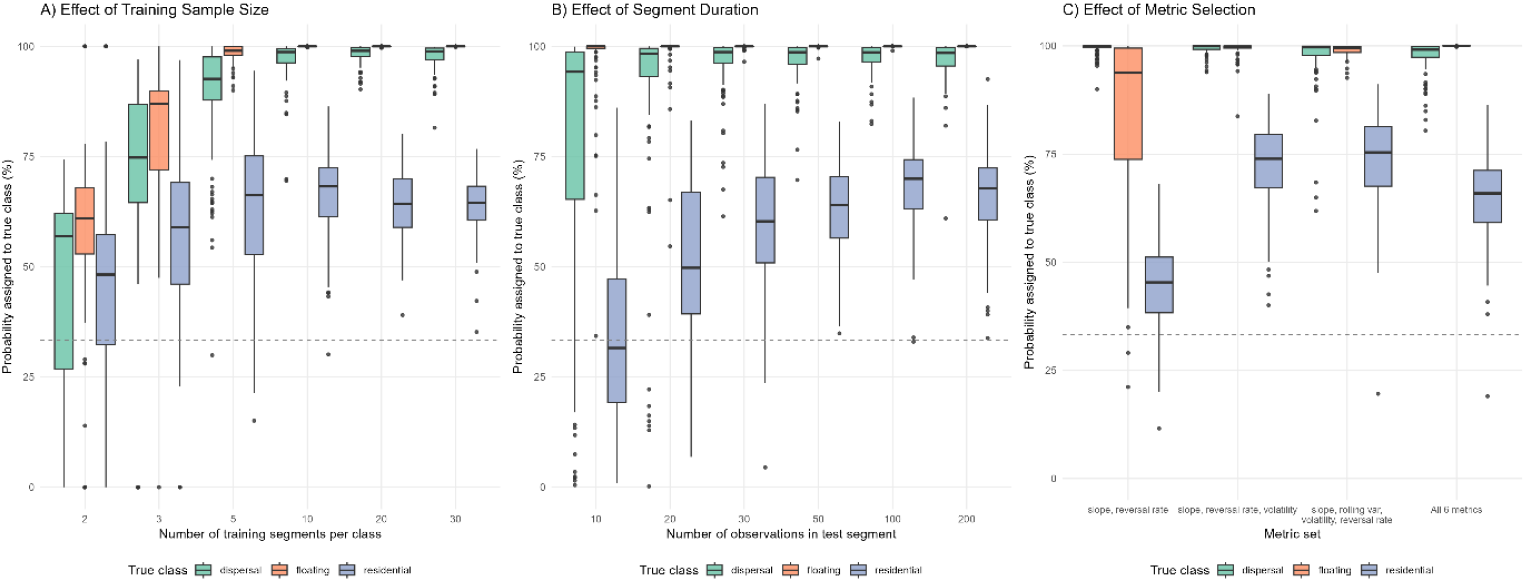
Effects of (a) training sample size, (b) segment duration, and (c) metric selection on classification accuracy. Dashed line indicates chance performance (33%). In (a), test segments contained 60 observations;in (b), 10 training segments per class were used;in (c), 10 training segments and 60-observation test segments. Boxplots show median and interquartile range across 100 replicates.

Using 10 training segments per class, segment duration similarly influenced classification confidence (Figure 3, panel B). Dispersal was reliably identified even with short segments of 10 observations, reflecting the strong directional signal captured by slope and autocorrelation metrics. In contrast, residential and floating classifications required longer observation periods to accumulate sufficient information for reliable discrimination. With 10 observations, residential segments received only 25–40%probability for the correct class, improving to 70–85%at 50 observations and approaching ceiling performance starting from 100 to more observations. This pattern reflects the inherent information content of different behavioral strategies: dispersal generates a distinctive monotonic NSD increase detectable within days, while distinguishing residential site fidelity from floating exploration requires observing patterns of return, reversal, and variance accumulation over weeks or months.

With 10 training segments per class and 60-observation test segments, metric selection affected classification performance differentially across behavioral categories (Figure 3, panel C). A minimal set comprising only slope and reversal rate achieved high accuracy for dispersal (median >95%) but performed poorly for residential classification (median ~45%), little better than by chance. Adding volatility as a third metric improved residential classification to approximately 65%, and the recommended four-metric combination (slope, rolling variance, volatility, reversal rate) performed comparably to the full six-metric (slope, residual standard error, rolling variance, volatility, reversal rate, autocorrelation) set across all strategies. This indicates that parsimonious metric selection, guided by exploratory analysis of training data distributions, can achieve robust classification without exhaustive metric computation. However, the specific metrics required depend on the behavioral categories being distinguished and the ecology of the species: slope alone suffices for dispersal detection, but residential-floating discrimination requires variance-based metrics capturing the characteristic instability of floating movement.

The confusion matrix aggregated across all training sizes revealed that classification errors were asymmetrically distributed among behavioral categories (Figure 4). Overall classification accuracy was 91.1%(1,636 of 1,796 trials correctly classified). Floating achieved the highest per-class accuracy at 97.5%, with misclassifications almost exclusively toward residency (2.5%) rather than dispersers (0%). Dispersal accuracy was 93.7%, with the 6.2%error rate consisting entirely of floating misclassifications, likely attributable to simulated stopovers creating temporary high-variance plateaus resembling floating behavior. Residential accuracy was lowest at 82.1%, with 17.7%of residential segments misclassified as floating and only 0.2%as dispersal. This asymmetric error pattern has a clear mechanistic interpretation: the excursions incorporated into residential trajectories create periods of elevated displacement and variance that, when sampled in isolation, resemble the erratic movement characteristic of floating behavior. The near-zero confusion between residential and dispersal reflects their fundamentally different NSD dynamics, oscillatory versus monotonically increasing which the slope and trend metrics readily distinguish.

**Figure 4.**
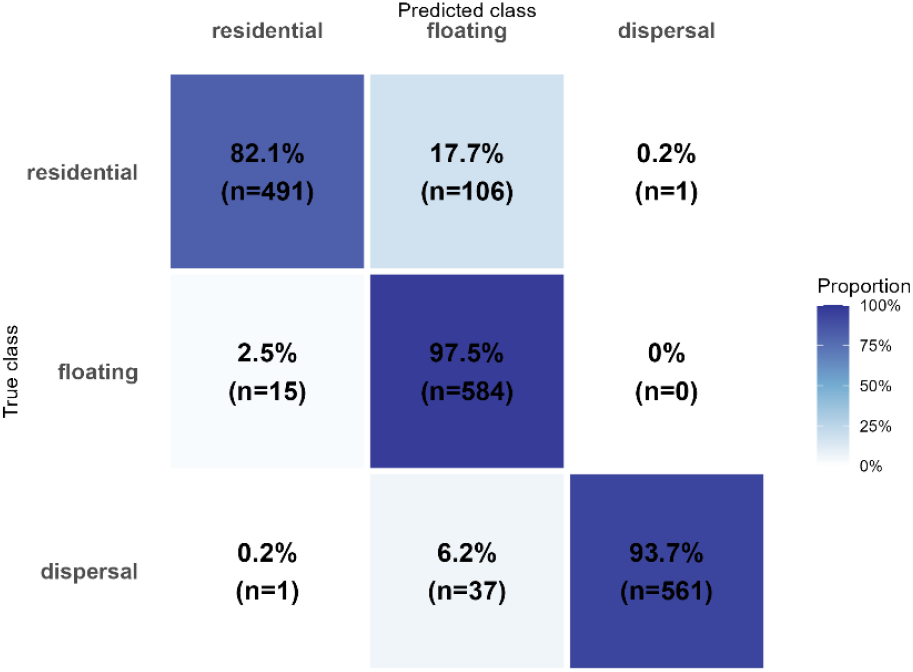
Confusion matrix for simulated trajectory classification (n = 1,796 trials). Values show percentage (count) of segments from each true class (rows) assigned to predicted classes (columns). Overall accuracy was 91.1%, with highest per-class accuracy for floating (97.5%) and lowest for residential (82.1%).

### 3.2 Case study 1: Gray Wolf

#### 3.2.1 Study species and movement strategies

Wolves exhibit three distinct movement strategies with important implications for population dynamics, territory establishment, and management. Territorial residents maintain stable home ranges within pack territories, typically characterized by low net displacement from a central location and consistent space use over extended periods.

Floating individuals engage in nomadic exploration without permanent settlement, representing wandering movements, after exclusion as pack members, or individuals searching for mating opportunities;these movements show irregular displacement patterns with frequent directional changes. Dispersal involves directed long-distance movements from (usually) natal territories to establish their new ones or intending to join the existing, characterized by sustained directional displacement.

#### 3.2.2 Data collection and training data

GPS telemetry data were collected from wolves across Central Europe (Czech Republic, Germany, Austria, and Poland) between 2016 and 2025. Training segments were identified through expert classification based on visual inspection of movement trajectories and knowledge of individual life histories. The training dataset comprised 18 segments from 13 individuals (Table 1).

**Table 1.** Training data composition for wolf behavioural classification, showing segment counts and selection criteria for each movement strategy.

| Behavioural class | N segments | Selection criteria |
| --- | --- | --- |
| Residential | 11 | Established territory holders with stable home ranges |
| Floating | 3 | Exploratory movements without site fidelity or directional persistence |
| Dispersal | 4 | Directed long-distance movements with clear start/end |

#### 3.2.3 Behavioural profiles

Six movement metrics were calculated for each training segment: slope, R^2^, rolling variance, volatility, reversal rate, and autocorrelation. The resulting behavioural profiles showed clear separation between movement strategies across all calculated metrics (Table 2). Residential behaviour was characterized by near-zero slopes indicating minimal net displacement, low R^2^ reflecting absence of directional trends, and high reversal rates typical of oscillatory movements within a confined home range. Dispersal showed the opposite pattern: Dispersal showed the opposite pattern, with high positive slopes, high R^2^, low reversal rates, and high autocorrelation reflecting persistent directional movement. Floating occupied an intermediate position with moderate slope values, high rolling variance reflecting spatial instability, and intermediate reversal rates characteristic of erratic, non-directional exploration.

**Table 2.** Behavioural profiles (mean ±SD) for wolf movement strategies. Metrics calculated from training segments (n = 11 residential, 3 floating, 4 dispersal).

| Metric | Residential | Floating | Dispersal |
| --- | --- | --- | --- |
| Slope | $-0.03 \pm 0.19$ | $1.89 \pm 3.44$ | $3629.02 \pm 2955.36$ |
| $R^2$ | $0.04 \pm 0.04$ | $0.04 \pm 0.02$ | $0.88 \pm 0.05$ |
| Rolling variance | $3.24e^4 \pm 9.97e^4$ | $5.47e^5 \pm 5.85e^5$ | $2.63e^8 \pm 3.66e^8$ |
| Volatility | $70.47 \pm 135.44$ | $279.24 \pm 48.95$ | $4543.49 \pm 4287.10$ |
| Reversal rate | $0.61 \pm 0.06$ | $0.48 \pm 0.07$ | $0.28 \pm 0.08$ |
| Autocorrelation | $0.32 \pm 0.23$ | $0.72 \pm 0.13$ | $0.89 \pm 0.03$ |

#### 3.2.4 Classification

We applied the classification framework to 30 wolves comprising 56 identified segments. The method assigned unambiguous classifications to 51 segments (91.2%), defined as segments where the highest-class probability exceeded 50%. The remaining 5 segments (8.8%) showed no class probability exceeding 70%and were flagged as ambiguous (Appendix 1). The distribution of classified phases reflects the demographic composition of the studied population, with residential individuals predominating and smaller proportions engaged in floating or dispersal behaviour: 30 residential (53.6%), 11 floating (19.6%), and 10 dispersal (17.9%) phases. As an illustrative example, wolf CZE15 displayed a 328-day tracking record with two breakpoints partitioning the trajectory into three behavioural phases (Figure 5, Table 3). Segment 1 (183 days, October 2024 –April 2025) was classified as residential with high confidence (92.8%, 95%CI: 89.7–95.8%), consistent with established territorial behaviour visible as stable, low NSD values in the trajectory. Segment 2 (37 days, April –May 2025) represented a dispersal event (97.8%, 95%CI: 82.5–100.0%), reflecting directed long-distance movement visible as a sharp NSD increase from near-zero to approximately 9,000 km^2^. Segment 3 (107 days, May –September 2025) was classified as floating (99.7%, 95%CI: 99.2–99.9%). This sequence represents a typical life-history transition from natal territory through dispersal to settlement-seeking behaviour.

**Table 3.** Classification results for wolf CZE15 showing bootstrap probability estimates (mean [95%CI]) for each behavioural class across three trajectory segments. Non-overlapping confidence intervals between classes indicate unambiguous classification. Overlapping intervals warrant additional scrutiny;however, if the confidence interval of the assigned class does not contain the mean probability of any alternative strategy, the classification remains reliable. If the confidence interval of the assigned class contains the mean of an alternative strategy, the classification should be treated with caution and ideally verified by expert review.

| Segment | Period | Z<sub>Residency</sub> | Z<sub>Floating</sub> | Z<sub>Dispersal</sub> | Class |
| --- | --- | --- | --- | --- | --- |
| 1 | 10 Oct–23 Apr | 92.8 [89.7, 95.8] | 7.1 [4.2, 10.3] | 0.0 [0.0, 0.1] | residency |
| 2 | 24 Apr–30 May | 0.0 [0.0, 0.0] | 2.2 [0.0, 17.5] | 97.8 [82.5, 100.] | dispersal |
| 3 | 31 May–14 Sep | 0.0 [0.0, 0.0] | 99.7 [99.2, 99.9] | 0.3 [0.1, 0.8] | floating |

**Figure 5:**
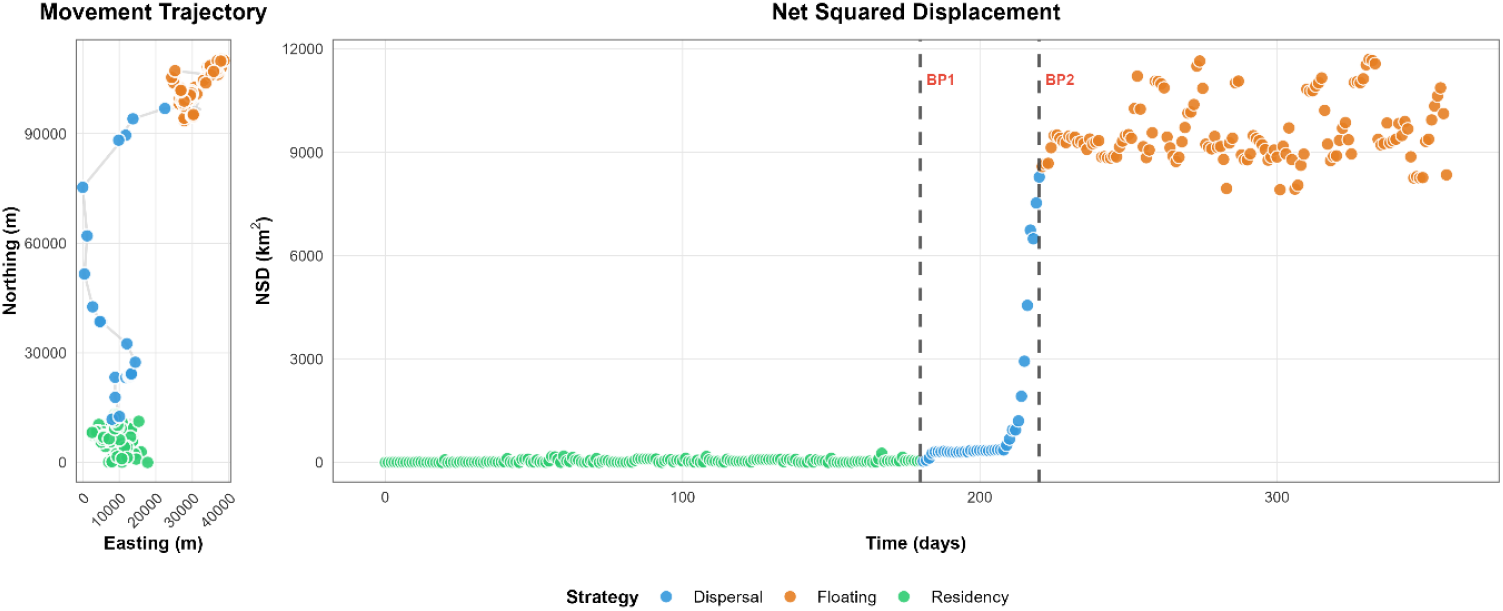
Movement trajectory and NSD time series for wolf CZE15. Left panel shows spatial positions coloured by classified movement strategy. Right panel displays NSD over time with breakpoints (BP1, BP2) separating three movement strategies: residency (green), dispersal (blue), and floating (orange).

### 3.3 Northern Lapwing

#### 3.3.1 Study species and movement strategies

Northern lapwings exhibit three distinct movement strategies that vary seasonally and across life-history stages. We defined three movement strategies based on expert classification. Residential phases encompass territory establishment, incubation, and chick-rearing with restricted movements around nest sites. Floating phases represent exploratory movements and migratory stopovers without directional persistence. Migration phases involve rapid, directed long-distance displacement between breeding grounds in Central Europe and wintering grounds in Spain and France. Distinguishing residential phases from floating behaviour proves particularly challenging, as both involve localised movements without sustained directional displacement (Eichhorn et al., 2017).

#### 3.3.2 Data collection and training data

GPS tracking data were collected from lapwings breeding in the Czech Republic and migrating to wintering grounds in Spain and France between 2024 and 2026. Positions were aggregated into 12-hour centroids to capture the faster behavioural transitions typical of migratory birds. Training segments were identified through expert classification based on visual inspection of NSD trajectories and knowledge of breeding phenology. The training dataset comprised 47 segments from four individuals, with all four individuals contributing segments to each behavioural class (Table 4), ensuring that profiles were not dominated by individual-specific movement patterns.

**Table 4.**
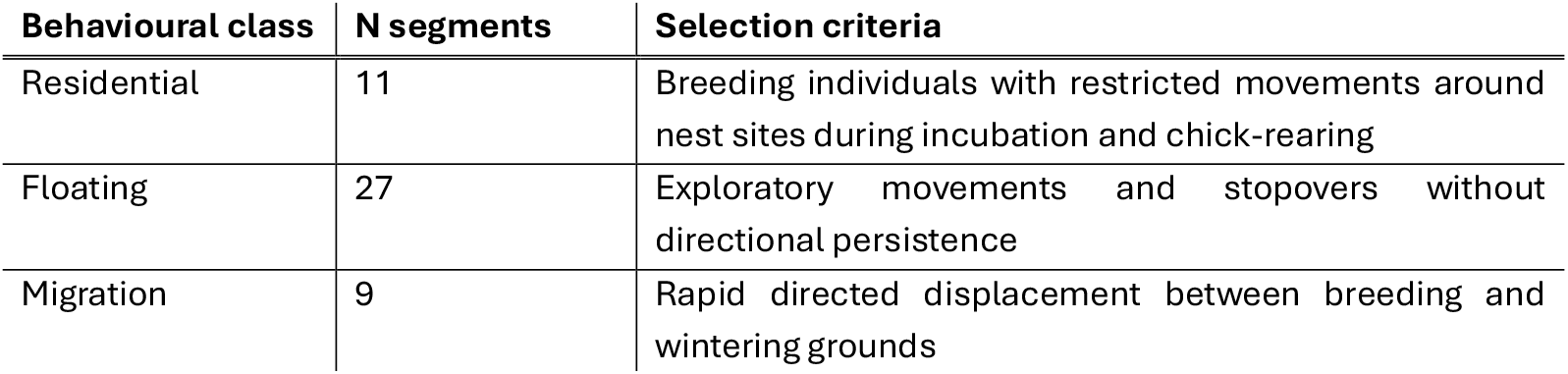
Training data composition for lapwing behavioural classification, showing segment counts and selection criteria for each movement strategy.

| Behavioural class | N segments | Selection criteria |
| --- | --- | --- |
| Residential | 11 | Breeding individuals with restricted movements around nest sites during incubation and chick-rearing |
| Floating | 27 | Exploratory movements and stopovers without directional persistence |
| Migration | 9 | Rapid directed displacement between breeding and wintering grounds |

#### 3.3.3 Behavioural profiles

Six movement metrics were calculated for each training segment: slope, R^2^, rolling variance, volatility, autocorrelation, and reversal rate. The resulting behavioural profiles showed clear separation for migration but considerable overlap between residential and floating strategies (Table 5). Migration was characterised by high slope values, high volatility, low reversal rates, and high R^2^, reflecting sustained directional displacement.

**Table 5.**
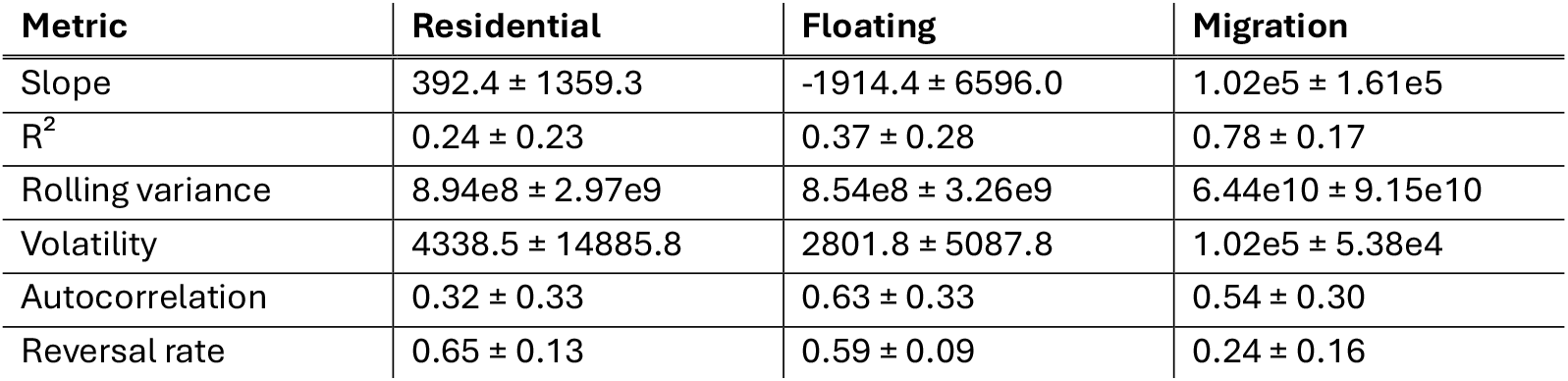
Behavioural profiles (mean ±SD) for northern lapwing movement strategies (n = 11 residential, 27 floating, 9 migration).

Residential and floating phases showed similar metric values across most dimensions, consistent with the genuine ecological similarity between breeding site fidelity and stopover behaviour (Table 5).

#### 3.3.4 Classification

We applied the classification framework to 90 segments across 11 individuals. Migration events were identified with high confidence, frequently achieving 100%probability with narrow confidence intervals. In contrast, residential and floating classifications typically showed probability distributions of approximately 50–55%versus 40–45%, with overlapping confidence intervals reflecting genuine ecological overlap between these strategies (Appendix 2). Classification confidence for residential and floating segments could be improved with larger training datasets, though the current uncertainty estimates honestly communicate the limits of available training data. Short-duration segments exhibited substantially wider bootstrap confidence intervals, a diagnostic signal indicating when additional scrutiny is warranted. Experts familiar with lapwing ecology confirmed that distinguishing breeding residency from post-breeding floating represents a genuine challenge, consistent with why most movement studies in this species focus on the migration-residency dichotomy rather than finer classification (Eichhorn et al., 2017). Our framework demonstrates that three-way classification is achievable, while bootstrap uncertainty quantification transparently identifies cases where confidence is limited.

As an illustrative example, individual 1300001473 displayed a 222-day tracking record partitioned into eight behavioural segments (Figure 6, Table 6). The trajectory captured the complete annual movement cycle of a migratory lapwing, beginning with a residential phase during the breeding season (1 May–16 June), followed by an extended post-breeding floating phase (16 June–19 September). Autumn migration proceeded in three discrete bouts separated by floating stopovers of varying duration, ending with wintering-ground floating behaviour (18 November–9 December). Migration bouts were classified with high confidence, while residential and floating phases showed characteristic probability overlap. Notably, despite this overlap, the 95%confidence intervals of the assigned class never contained the mean probability of any alternative movement strategy, supporting the reliability of all classifications. Segment 7 (12–18 November, 6 days) nonetheless exhibited notably wide confidence intervals for both migration [20.1, 100.0] and floating [0.0, 79.9], reflecting high classification uncertainty driven by the short segment duration, a diagnostic signal that this segment warrants additional scrutiny.

**Figure 6:**
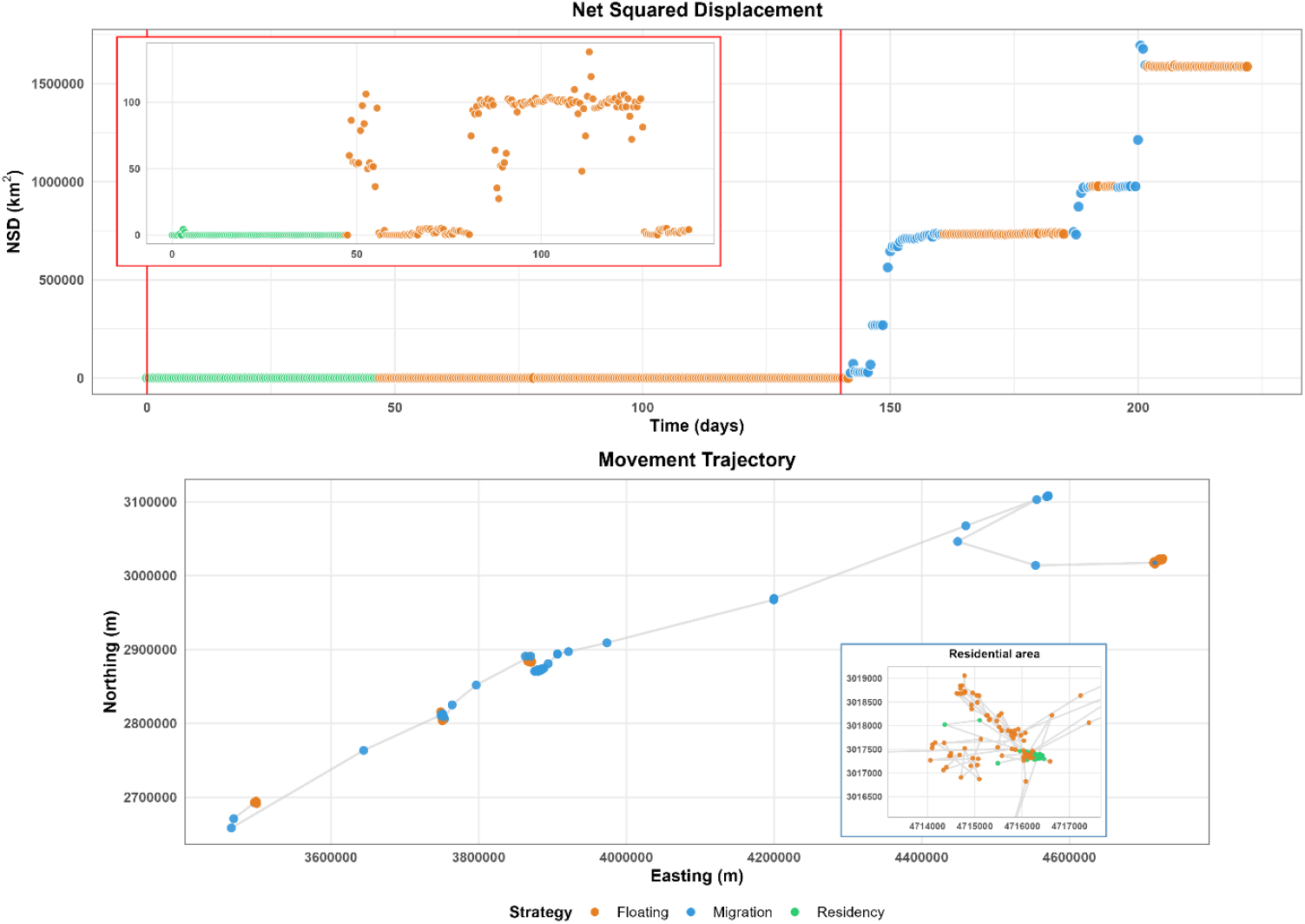
Movement trajectory and NSD time series for lapwing (ID: 130000147). Lower panel shows spatial positions coloured by classified movement strategies. Upper panel displays NSD over time with three movement strategies. : residency (green), migration (blue), and floating (orange).

**Table 6.**
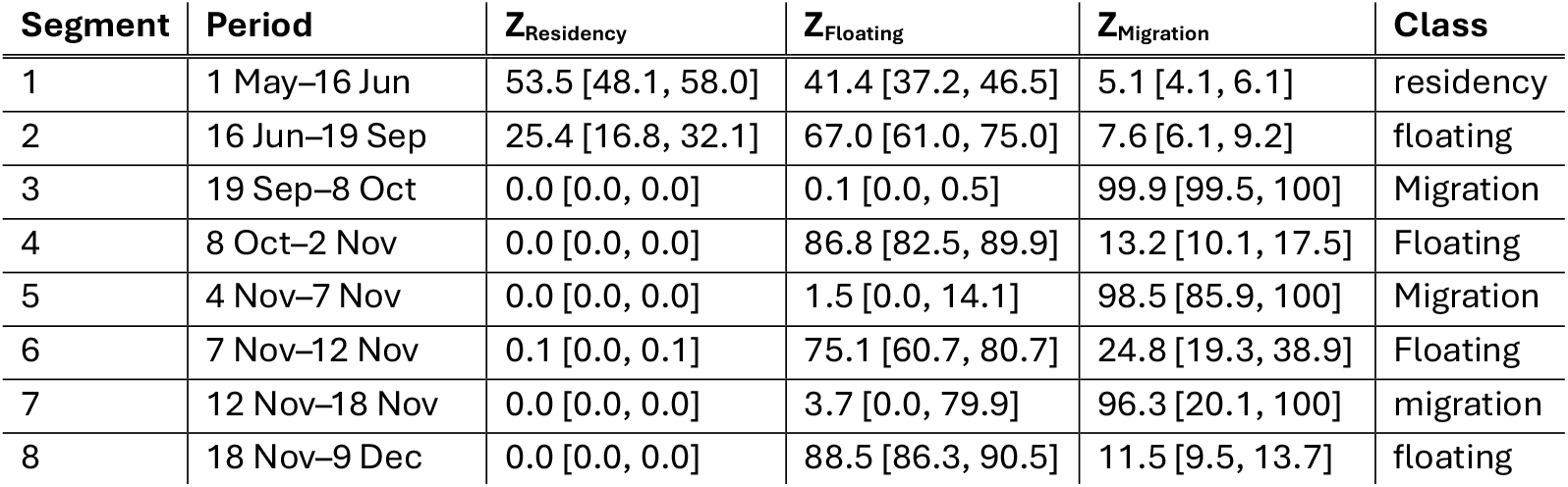
Classification results for lapwing (ID: 130000147) showing bootstrap probability estimates (mean [95%CI]) for each movement strategies across eight trajectory segments. Non-overlapping confidence intervals between classes indicate unambiguous classification. Overlapping intervals warrant additional scrutiny;however, if the confidence interval of the assigned class does not contain the mean probability of any alternative strategy, the classification remains reliable. If the confidence interval of the assigned class contains the mean of an alternative strategy, the classification should be treated with caution and ideally verified by expert review.

## 4 Discussion

Classifying animal movement strategies from tracking data remains a challenge in movement ecology. Existing approaches often require either strong parametric assumptions about trajectory shape or extensive labelled datasets for machine learning (Bunnefeld et al., 2011;Valletta et al., 2017), creating barriers for researchers working with novel species or limited sample sizes or without extensive knowledge of statistics, since telemetry is often used outside the professional scientific community. The profile-based framework presented here offers an alternative option: empirically derived behavioural profiles, distance-based classification, and bootstrap uncertainty quantification that together provide reliable, interpretable results while remaining accessible to ecologists without specialized statistical training.

The simulation study provideds a foundation for practical applications. A central question facing any researcher wishing to use our framework is: how much training data do I need? Our results suggest classification accuracy improved dramatically between 2 and 5 training segments per behavioural class, then largely stabilized. With 10 segments per class, dispersal and floating classifications did not improve further while residential accuracy reached approximately 80%. Additional training data beyond this threshold yielded diminishing returns. This finding has an important practical implication: researchers do not need to wait with classifications until they collected an extensive training dataset. A carefully curated data set of 5 to 10 unambiguous examples for each movement strategy, selected based on expert knowledge or independent validation, provides sufficient foundation for reliable classification.

Segment duration influences classification confidence in ways that reflect the underlying information content of different movement strategies. Dispersal was reliably identified even from short segments of only 10 observations, because constant directional movement results in a distinctive NSD signature that slope and autocorrelation metrics can capture almost immediately. Residential and floating behaviours, by contrast, require longer observation periods as heir discrimination relies on the accumulation of characteristic oscillatory NSD patterns (repeated reversals and local variance) that only become statistically reliable over weeks of observation.. Our simulations showed residential classification improving substantially between 10 and 50 observations and approaching reliable performance at 100 or more observations. For large mammals tracked at daily temporal resolution, this translates to minimum recommended segment durations of 30 to 60 days. Shorter segments can still be used, but researchers should expect wider confidence intervals honestly reflecting the limited information available.

The confusion matrix from simulation experiments suggested an asymmetric error structure that illuminates fundamental similarities and differences among movement strategies. Residential segments were misclassified as floating in 17.7%of trials, while the reverse error occurred in only 2.5%of cases. This asymmetry makes mechanistic sense: residential trajectories incorporate excursions representing hunting forays, territorial patrols, or exploratory movements that, when sampled in isolation, resemble the erratic displacement patterns characteristic of floating behaviour. Floating, however, rarely mimics the consistent return-to-centre pattern defining residency. Dispersal was confused with neither residential (0.2%) nor floating confused with dispersal (0%), reflecting the fundamentally different NSD dynamics of sustained directional movement. The 6.2%dispersal-to-floating misclassification rate likely reflects simulated stopovers creating temporary high-variance plateaus. These patterns from simulated data strikingly parallel our empirical observations: wolves showed clean separation between residential and dispersal, while lapwings exhibited substantial residential-floating overlap that experts confirmed reflects genuine ecological ambiguity rather than methodological failure.

The framework differs from existing approaches in three key aspects: flexibility in handling diverse trajectory shapes, interpretability of classification decisions, and modest training data requirements. Parametric NSD model fitting assumes trajectories conform to predefined functional forms, typically asymptotic curves for migration or dispersal. This assumption fails when animals exhibit complex or variable movement patterns, and fitting algorithms often struggle with the substantial NSD variance characteristic of floating behaviour. Our profile-based approach imposes no functional form, instead learning characteristic metric distributions directly from training data (Börger & Fryxell, 2012;Bunnefeld et al., 2011;Spitz et al., 2017). Hidden Markov models offer sophisticated tools for movement strategy inference but require specifying state transition probabilities and emission distributions, assumptions that may be difficult to justify for species with limited prior behavioural characterization(Cisneros-Araujo et al., 2026;McClintock & Michelot, 2018;Michelot et al., 2016). Machine learning methods can achieve high classification accuracy but typically require larger training datasets than movement ecologists commonly possess, and their black-box nature complicates ecological interpretation (Valletta et al., 2017;Wijeyakulasuriya et al., 2020). The profile-based framework occupies a useful middle ground: more flexible than parametric fitting, more interpretable than machine learning, and less assumption-heavy than HMMs.

The cross-taxa case studies demonstrate that this flexibility translates into practical utility. Wolf and lapwing movement occur on fundamentally different scales: wolf dispersal slopes averaged 3,629 km^2^ per day while lapwing migration slopes reached 102,000 km^2^ per day, a nearly 30-fold difference. Yet the same methodological framework, parameterized with species-appropriate training data, achieved reliable classification in both systems (Abrahms et al., 2017). This adaptability reflects a key design principle: rather than hard-coding thresholds or functional forms, the framework learns what distinguishes movement strategies in each specific context. A slope of 50 km^2^ per day might indicate dispersal in a territorial songbird but residency in a wide-ranging carnivore;the profile-based approach handles both cases by referencing empirical training distributions rather than fixed values.

The bootstrap confidence intervals, while a standard tool for uncertainty quantification, serve as an additional diagnostic function in this framework. Wide confidence intervals for short segments do not indicate methodological failure;they honestly acknowledge that brief observation periods contain insufficient information for confident classification. Similarly, overlapping probabilities for behaviourally similar strategies (residential versus floating in lapwings) reflect genuine ecological ambiguity. When experts themselves struggle to distinguish breeding-site fidelity from localized post-breeding exploration based on movement patterns alone, a method that reports 55%residential versus 42%floating probability is providing valuable information: this trajectory does not clearly belong to either category. Researchers can use such signals to flag segments requiring additional information (e.g., reproductive status, proximity to nest sites) or to acknowledge uncertainty in downstream analyses rather than propagating false confidence.

### Practical recommendations

Based on our simulation experiments and empirical experience, we offer the following guidelines for researchers implementing this framework. A key advantage of this framework is reliable classification with relatively small training samples, compared to hundreds typically required for machine learning approaches, though the minimal sample size depends on the complexity of the behaviours and their degree of separation in metric space. First, assemble training data comprising at least 5 to 10 segments per movement strategy, prioritizing unambiguous examples over quantity. Segments selected for training should represent clear instances of target behaviours based on expert knowledge, independent observation, or life-history information. Second, examine training data distributions before finalizing metric selection. Not all metrics contribute equally across all classification tasks: slope and autocorrelation dominate dispersal detection, while variance-based metrics prove essential for distinguishing residential from floating behaviour. Metrics showing complete overlap between categories add noise without information and should be excluded. Third, apply minimum segment duration thresholds appropriate to your study system. For large mammals with daily fixes, we recommend minimum durations of 30 to 60 days;for species with faster behavioural transitions (e.g., breeding birds), shorter segments may suffice but will produce wider confidence intervals. Fourth, interpret classification probabilities as meaningful signals rather than nuisance uncertainty. Segments with ambiguous classifications often correspond to transitional periods, mixed behaviours, or genuinely intermediate movement patterns. Finally, for datasets with small training samples (fewer than 15 to 20 segments total), consider single rather than double bootstrap to avoid instability from resampling already-limited data.

Several limitations warrant acknowledgment. Classification quality depends entirely on training data quality;unrepresentative or mislabelled training segments will produce unreliable profiles. The framework assumes identified segments represent relatively homogeneous movement patterns, an assumption that may be violated during rapid behavioural transitions. Optimal metric selection requires exploratory analysis and some degree of ecological judgment, introducing subjectivity into an otherwise algorithmic process. And while bootstrap confidence intervals provide useful uncertainty estimates, they do not account for all sources of error, particularly systematic biases in GPS positioning or temporal autocorrelation in location data.

Future development should address several promising directions: real-time classification for adaptive wildlife management, where incoming tracking data trigger alerts when animals transition between behavioural states;integration with step selection functions for state-dependent habitat analysis;and extension to three-dimensional movement data from marine or aerial tracking. An R package (moveprofile) implementing the complete framework is available at https://github.com/kadleci/moveprofile.

## 5 Conclusion

The profile-based framework presented here provides an interpretable, flexible, and uncertainty-aware approach to classifying animal movement strategies from tracking data. Three findings stand out. Reliable classification can be achieved with as few as 5 to 10 training segments per category, well within reach of typical telemetry studies. The same framework transfers across taxa with very different spatial and temporal scales, provided that training data are species appropriate. Bootstrap confidence intervals serve as diagnostic indicators of genuine ecological ambiguity rather than mere statistical noise, flagging cases where movement patterns themselves resist confident assignment.

We emphasize that this framework does not replace deeper mechanistic understanding of animal movement. It provides a standardized, reproducible tool for behavioural classification that can serve as a foundation for subsequent analyses: linking movement strategies to habitat selection, quantifying individual variation in life-history strategies, or identifying population-level patterns in space use. By making reliable classification accessible without requiring extensive statistical expertise, we hope to lower barriers for movement ecologists seeking to extract behavioural insight from increasingly abundant tracking data.

An R package implementing the complete workflow, from raw GPS coordinates through classified trajectories with uncertainty estimates, is available at https://github.com/kadleci/moveprofile and will be submitted to CRAN upon publication.

## 6 Acknowledgements

We would like to express our sincere gratitude to all contributors in the field, namely Jan Mokry, Lukas Zak, Vojtech Oldrich, Jan Hornicek, Andreas Berger, Michal Bojda, Christian Eder, Šárka Frýbová, Jan Koranda, JiříLabuda, Karolina Mikslováand Simon Zauner. We are thankful for the assistance of Hana Horáková, Vladimír Piaček and other veterinarians during wolf trapping. We would especially like to thank Ashley McLaren, Bryce Olson, Ilka Reinhardt and Steve K. Windels for sharing their experience, advice and training in wolf trapping. We are grateful to Šumava NP, Czech Switzerland NP, the Nature Conservation Agency of the Czech Republic (Beskydy PLA and Krkonoše NP Administrations), the Austrian Ministry of Defense, the Ministry of the Environment of the Czech Republic, and Lesy České republiky for their cooperation and support. We would like to thank Miroslav Šálek, Kateřina Trejbalová, Václav Zámečník, Michaela Kadavá and Petr Chajma with assistance during lapwing catching.

## Permits

Wolf trapping was authorised by local nature protection authorities and followed ethical standards set by collaborating universities and governments: Czech University of Life Sciences (MZP/2018/630/2582, MZP/2022/630/929, SZ NPS 08816/2020/4, SZ SNPCS 00118/2022, KUUK/049576/2022), Mendel University (SR/0012/US/2021-5, MZP/2022/630/735), Krkonoše NP Administration (01273/2021 and 09508/2023), Nature Conservation Agency of the Czech Republic (Implementation Contract from 16.8.2021), Ministry of the Environment (MZP/2022/630/735), and Government of Lower Austria (LF1-TVG-59/001-2018). Licences for the German part of the wolf research (Dresden University of Technology) were issued by the Animal Welfare Authority of the State of Mecklenburg-Western Pomerania (7221.3-1-1-056/11-1;7221.3-1-027/15;7221.3-1-046/20), the Animal Welfare Authority of the German Armed Forces (42-34-30-20 G02-24), and the Animal Welfare Authority of the State of Saxony-Anhalt (203.m-42502-2-1686 TUDD G). The capture of northern lapwings was carried out under a ringing licence issued by the ringing station of the National Museum in Prague (licence no. 1082);the attachment of telemetry devices was authorised by the municipal authorities of Hradec Králové(MMHK/254806/2024), Holice (MUHO 10216/2024/OŽP/Po), and Pardubice (SZ_MMP 55492/2024).

## 8 Statement on inclusion

This study was conducted across Central Europe, with GPS telemetry data collected in the Czech Republic, Germany, Austria and Poland. The authorship team includes researchers based in the countries where data were collected, representing institutions in the Czech Republic, Austria and Germany. Local collaborators and data providers, including national park administrations, nature conservation agencies and wildlife management authorities, were involved throughout the research and are acknowledged accordingly. The analytical framework was developed in close collaboration between researchers with direct field experience in the study region and methodological expertise, ensuring that ecological context informed all stages of method development and validation.

